# Within-individual changes reveal increasing social selectivity with age in rhesus macaques

**DOI:** 10.1101/2022.05.31.494118

**Authors:** Erin R. Siracusa, Josué E. Negron-Del Valle, Daniel Phillips, Michael L. Platt, James P. Higham, Noah Snyder-Mackler, Lauren J. N. Brent

## Abstract

Accumulating evidence in humans and other mammals suggests older individuals tend to have smaller social networks. Uncovering the cause of these declines is important as it can inform how changes in social relationships with age might affect health and fitness in later life. Smaller social networks might be detrimental, but may also be the result of greater selectivity in partner choice, reflecting an adaptive solution to physical or physiological limitations imposed by age. While greater selectivity with age has been shown in humans, the extent to which active ‘social selectivity’ within an individual’s lifetime occurs across the animal kingdom remains an open question. Using 8 years of longitudinal data from a population of free-ranging rhesus macaques we provide the first evidence in a non-human animal for within-individual increases in social selectivity with age. Going beyond previous cross-sectional studies, our within-individual analyses revealed that adult female macaques actively reduced the size of their networks as they aged and focused on partners previously linked to fitness benefits, including kin and partners to whom they were strongly and consistently connected earlier in life. Females spent similar amounts of time socializing as they aged, suggesting that network shrinkage does not result from lack of motivation or ability to engage. Furthermore, females remained attractive companions and were not isolated by withdrawal of social partners. Taken together, our results provide rare empirical evidence for social selectivity in non-humans, suggesting patterns of social aging in humans may be deeply rooted in primate evolution and may have adaptive value.

**Significance statement:** The narrowing of social networks and prioritization of meaningful relationships with age is commonly observed in humans. Determining whether social selectivity is exhibited by other animals remains critical to furthering our understanding of the evolution of late-life changes in sociality. Here we test key predictions from the social selectivity hypothesis and demonstrate that female rhesus macaques show within-individual changes in sociality with age that resemble the human social aging phenotype. Our use of longitudinal data to track how individuals change their social behavior within their lifetimes offers the most conclusive evidence to date that social selectivity is not a phenomenon unique to humans and therefore might have deeper evolutionary underpinnings.

## Introduction

Social relationships change in quality and quantity across the lifespan (1, 2), a phenomenon that has been referred to as ‘social aging’ (3, 4). Older people have commonly been observed to engage in less social activity and to have smaller social networks (1). Given the established health benefits of social integration (5, 6) this has led to an increasing concern of an ‘epidemic’ of social isolation among the elderly (7). However, research over the past several decades has suggested that reduced social network size may not simply be the result of unfavorable conditions in old age, such as increased frailty and reduced social competence (8). Instead, “socioemotional selectivity theory” proposes that aging individuals proactively focus on meaningful relationships, such as close friends and relatives, possibly as a result of their increased awareness of limited future time (9, 10).

Intriguingly, social selectivity in older individuals may not be limited to humans. Several species of non-human primates have been suggested to show patterns of social aging that might indicate greater selectivity in social partners in later life (3, 4, 11, 12). Whether or not non-human primates have future-time awareness (13), evidence for social selectivity in some of our closest living relatives could hint at deep evolutionary origins behind this phenomenon and may suggest that selectivity is an adaptive strategy that individuals use to cope with the physical and physiological limitations they face as they age.

Fundamentally, social selectivity is driven by within-individual changes in behavior with age. However, to date, most non-human studies have been cross-sectional in nature. That is, they have shown older animals differ from younger ones, but have either not had the longitudinal data needed to track changes within aging individuals or have not disentangled within-individual changes due to age from among-individual differences that might instead result from differences among cohorts or processes like selective disappearance (14, 15). For instance, if more social individuals are more likely to die because of increased exposure to disease or increased levels of competition, then an apparent age-related decline in sociality might appear at the population level without necessitating any within-individual change. In studies that are unable to separate within-from between-individual effects it is therefore not possible to conclusively demonstrate that social selectivity is driving the observed age-related patterns.

In addition to being necessarily driven by changes within aging individuals, the social selectivity hypothesis implies that individuals actively and adaptively narrow their networks with age. This makes it important to rule out alternative explanations for age-based reductions in sociality. For example, declines in social engagement might be driven by loss of social interest, motivation, or physical ability to engage (16). Alternatively, older individuals might be perceived as less valuable partners (e.g. due to declines in social status; 17, 18) resulting in reductions in network size as a result of withdrawal of social partners. Apparent preferences for related individuals with age might also reflect demographic changes and the loss of familiar, unrelated partners (such as age mates) due to mortality (19). Some progress has been made in generating evidence in favor of the occurrence of active social selectivity in non-humans (see 3, 4, 11, 12 for examples), but differentiating between these alternative explanations necessitates a clear set of predictions for what we would expect to see if social selectivity were actively occurring as an individual ages and the data to test these predictions.

In this study we used within-individual data from a longitudinal study of highly social rhesus macaques (*Macaca mulatta*) to test the four main behavioral predictions we derived from the sociality selectivity hypothesis: first that social networks narrow as an individual ages. That is, individuals interact with fewer social partners as they get older. Second, that this narrowing is driven by the aging individual in question rather than by the withdrawal of their social partners. Third, that aging individuals remain actively engaged with others. This prediction was intended to disentangle social selectivity from a narrowing of networks that might result from the loss of interest, motivation, or physical ability to engage socially. Fourth, we predicted that older individuals would focus their social effort on important partners. In the bio-gerontology literature “important” relationships have typically been viewed as family and close friends because they are the most emotionally salient relationships (20). Increased focus on kin and strongly-connected partners with age, however, might also make sense from an evolutionary perspective since these are partners that commonly provide important direct and indirect fitness benefits (21-23).

We tested these predictions using a long-term dataset spanning eight years and 204 individuals. Our subjects were mature adult females aged 10 to 28 (mean age = 14.3) from 6 naturally-formed mixed-sex social groups from the well-studied population living on the island of Cayo Santiago. We chose this age range because we were specifically interested in looking at age-based changes in social behavior from prime adulthood into old age. At six years old females are deemed adults (24) and analyses from the Cayo Santiago population have shown that the median lifespan for females that survive to reproductive age is 18 years, with a maximum lifespan of about 30 years (25, 26). For the 204 females that we monitored between 2010 and 2017 we had on average 2.8 years of data per individual with a range of 1 to 8 years (Fig. 1). We collected two different types of social interaction data - grooming interactions and spatial proximity – which we used in our analyses as measures of social connectedness, in line with many other studies on Cercopithecine primates (21, 27-30).

**Fig. 1.**
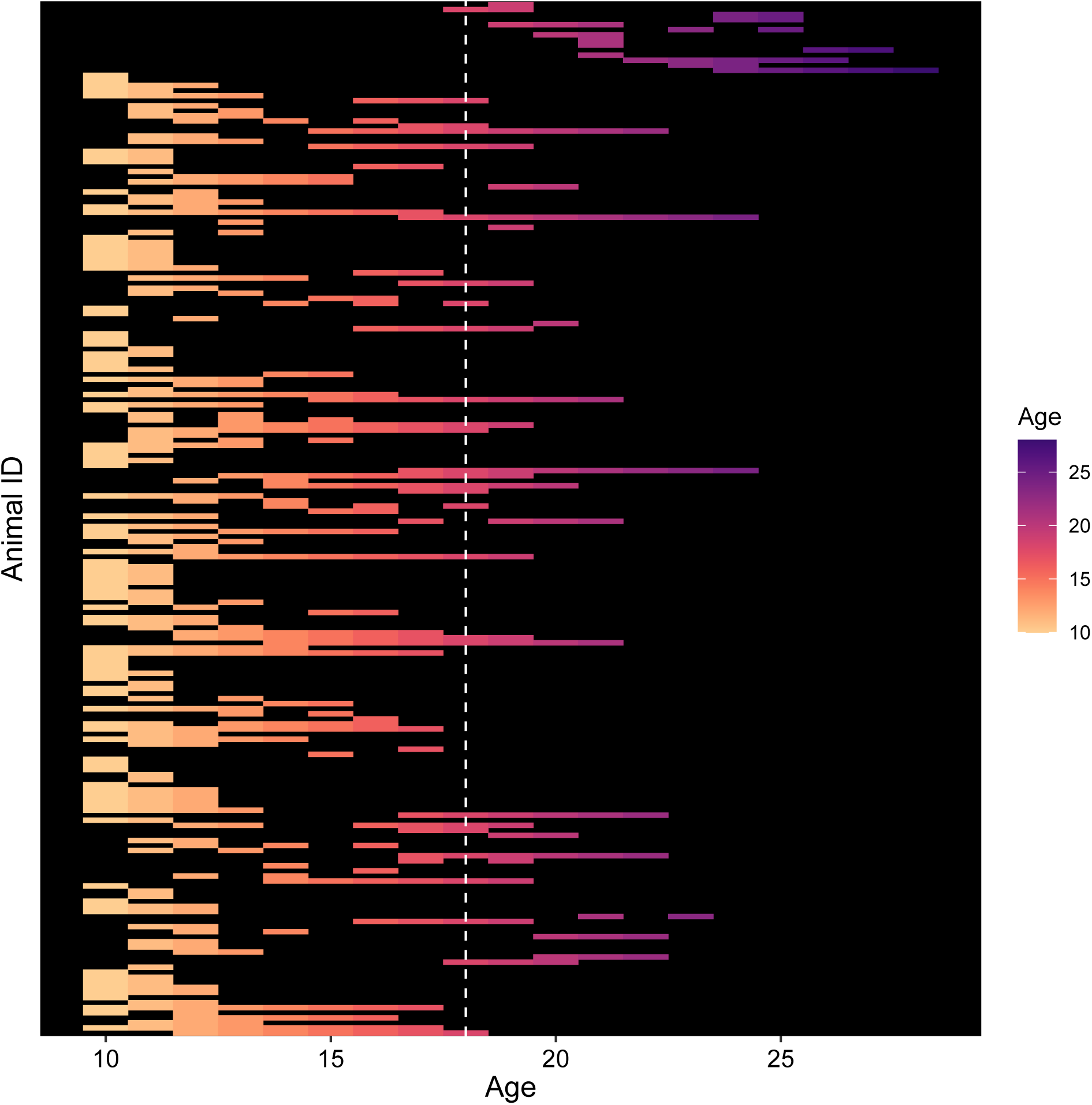
Heatmap showing age ranges over which each individual was sampled. Each row represents one individual. The dotted white line at age 18 indicates the median age of death in this population.

Crucially, to test that social selectivity was driven not by differences between individuals of different ages, but by changes in behavior within an individual as they grew older, we used a within-individual centering approach (15). We provide details of this approach in the Methods but briefly, within-individual centering is necessary when there is natural variation in the range of a given predictor variable (x) over which individuals are sampled (15). In this study, there was variation in the age ranges over which we had behavioral data for each female macaque. For example, we might have data from one female between the ages of 12 to 19 while for another female we have data from 20 to 27 (see Fig. 1). The inclusion of random intercepts in mixed models is not sufficient to account for this variation as such random effects can account for between-subject variation in the response variable y (i.e., a given metric of social behavior), but they do not automatically account for between-subject variation in x (i.e. age). In such situations, an association between x and y might be driven by a within-subject effect of x on y (individuals change their social behavior as they age) or by a between-subject effect of x on y (individuals with high average age also have high average sociality) (15).

To distinguish between these two alternative explanations we fitted models where we separated age into two terms: the between-subject effect was obtained by taking the average age of each subject (called “average age” in the models) and the within-subject effect was calculated by subtracting average age from each age at which the individual was observed (called “within-individual age” in the models). This “within-individual age” term is therefore the primary term of interest and represents how an individual’s deviation from its mean age affects its behavior. Additionally, because female rhesus macaques have a strict dominance hierarchy whereby daughters occupy ranks immediately below their mother (31), we tested for an interaction between rank and within-individual age in all of our models to assess whether social status affected how individuals changed their social behavior as they aged (19, 32). Finally, we fitted within-individual age as a random slope term over individual ID in all models to assess if there was among-individual variation in how females changed their social behavior with age. We present all effect sizes below by describing the expected within-individual change in social behavior over the study period (8 years) for a mid-ranking individual while holding average age constant at the mean (14.3 years). Our results provide the first evidence in a non-human animal for within-individual increases in social selectivity with age.

## Results

### Prediction 1: Social networks narrow with age

Female rhesus macaques showed within-individual declines in their number of social partners with age. On average, females reduced their number of grooming partners by 44% over an eight-year period (within-individual age: *β* = -0.06; 95% CI = -0.12, -0.01; Figure 2a,b, Table S1) and also reduced the number of partners that they spent time in proximity to, although this was modulated by dominance rank (Figure 2c,d; Table S2). Relative to their mean age, low-ranking individuals reduced their number of proximity partners by 47% over an eight-year period while high-ranking individuals only reduced their number of proximity partners by 8% (within-individual age:rankL: *β* = -0.07; 95% CI = -0.13, -0.01).

**Fig. 2.**
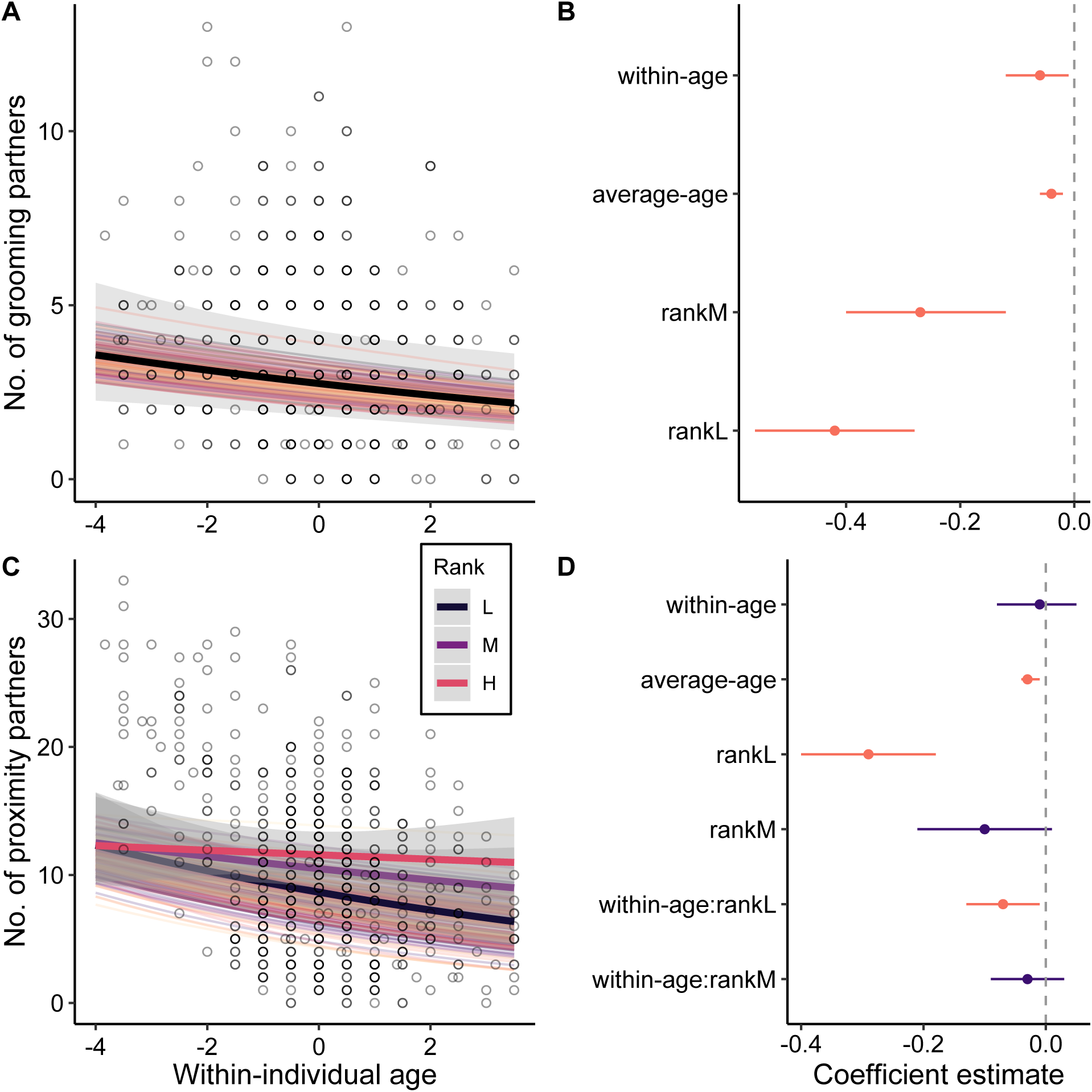
Narrowing of social networks with age. (**A** and **C**) Females show within-individual declines in number of (**A**) grooming partners and (**C**) proximity partners with age. Points represent raw data with the thick solid lines showing the average predicted within-individual change (dependent on rank where appropriate). Random slopes are shown using the thin coloured lines to illustrate the amount of inter-individual variation in a given social-aging pattern. Shaded gray bars indicate 95% confidence intervals around the predicted values. (**B** and **D**) Effect sizes and 95% credible intervals for all fixed effects and interaction terms for models that test the effect of age on the number of (**B**) grooming partners and (**D**) proximity partners. Instances where the 95% CI overlaps zero are colored in purple.

### Prediction 2: Narrowing of social networks is driven by the aging individual

Female macaques showed a within-individual decrease in the number of partners that they approached with age, which was modulated by rank (Figure 3a,b; Table S3). Relative to their mean age, low-ranking individuals reduced the number of individuals they approached by 51% over an eight-year period while high-ranking individuals reduced the number of individuals they approached by 15% (within-individual age:rankL: *β* = -0.08; 95% CI = -0.14, -0.01). However, females continued to be approached by similar numbers of partners as they aged regardless of rank (within-individual age: *β* = -0.02; 95% CI = -0.07, 0.02; Figure 3c,d, Table S4). Therefore, the narrowing of networks with age appears to be driven by within-individual behavioral changes of the aging individual rather than the withdrawal of social partners.

**Fig. 3.**
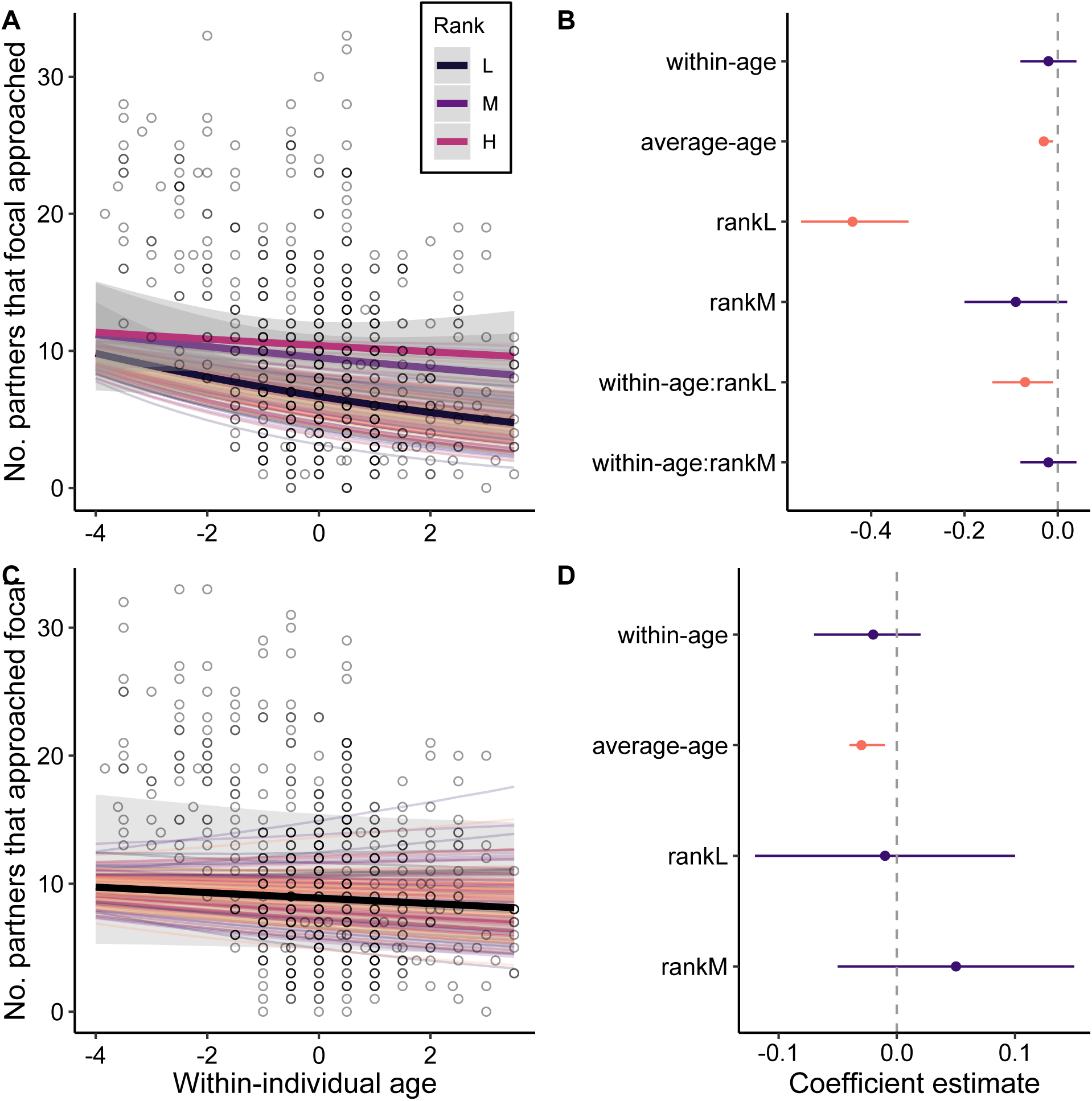
Network narrowing is driven by the aging individual. (**A** and **C**) Females show (**A**) within-individual declines in the number of partners that they approached, but (**C**) no change in the number of partners that they are approached by with age. Points represent raw data with the thick solid lines showing the average predicted within-individual change (dependent on rank where appropriate). Random slopes are shown using the thin coloured lines to illustrate the amount of inter-individual variation in a given social-aging pattern. Shaded gray bars indicate 95% confidence intervals around the predicted values. (**B** and **D**) Effect sizes and 95% credible intervals for all fixed effects and interaction terms for models that test the effect of age on (**B**) number of partners approached and (**D**) number of partners approached by. Instances where the 95% CI overlaps zero are colored in purple.

### Prediction 3: Individuals show continued engagement and interest in the social world as they age

Female macaques continued to give (within-individual age: *β* = -0.05; 95% CI = -0.12, 0.03; Figure 4a,b; Table S5) and receive (within-individual age: *β* = 0.02; 95% CI = -0.05, 0.09; Figure 4c,d; Table S6) similar amounts of grooming regardless of their age. Similarly, females did not show a change in the amount of time they spent in proximity to other individuals with age (within-individual age: *β* = -0.02; 95% CI = -0.08, 0.05; Figure 4e,f; Table S7). Therefore, while females interacted with fewer partners as they aged, they continued to spend similar amounts of time on social behavior, suggesting they remained motivated and engaged in the social world.

**Fig. 4.**
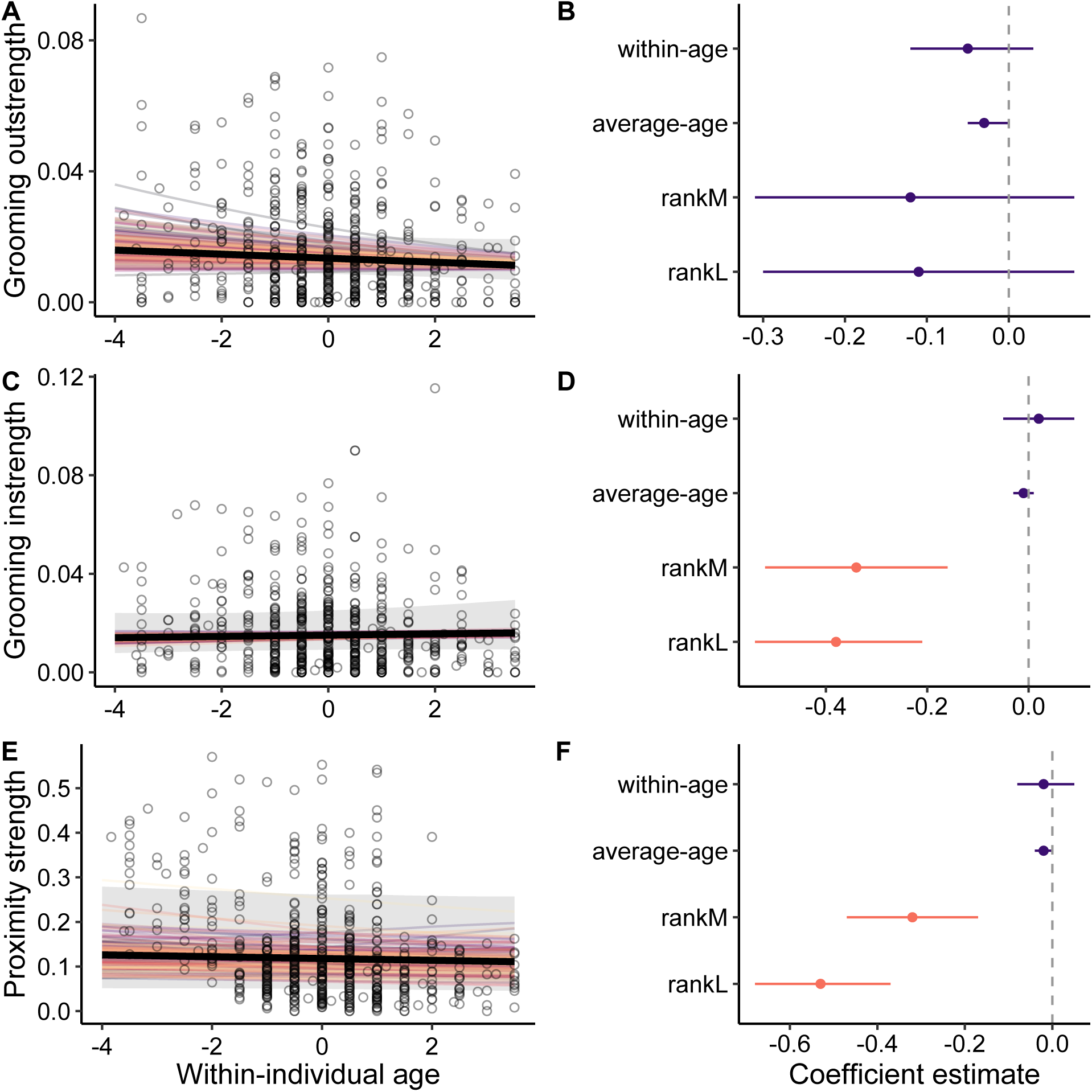
Individuals show continued engagement and interest in the social world as they age. (**A, C** and **E**) Females show no within-individual change in the amount of time spent (**A**) giving and (**C**) receiving grooming and (**E**) in proximity to other females as they age. Points represent raw data with the thick solid lines showing the average predicted within-individual change. Random slopes are shown using the thin coloured lines to illustrate the amount of inter-individual variation in a given social-aging pattern. Shaded gray bars indicate 95% confidence intervals around the predicted values. (**B, D** and **F**) Effect sizes and 95% credible intervals for all fixed effects for models that test the effect of age on amount of time spent (**B**) giving and (**D**) receiving grooming and (**F**) in proximity to other females. Instances where the 95% CI overlaps zero are colored in purple.

### Prediction 4: Individuals focus on important relationships in later life

Previous research on rhesus macaques has shown that females preferentially form relationships with female kin (33, 34) and that females with strong and stable connections to favored social partners have significantly reduced mortality risk (21). It therefore seems likely that both **(a)** kin, **(b)** strongly connected partners and **(c)** stable partners may all be important social relationships that females might strive to maintain in later life. To test whether females focused on important relationships as they aged we quantified changes in these three types of “important” partners. We assessed first whether **(a)** females increased their proportion of kin partners with age. Second, on a dyadic rather than individual level, we assessed whether, at their last time point in the dataset (i.e. oldest age), females were more likely to choose partners **(b)** to whom they had previously been strongly connected or **(c)** with whom they previously had stable connections. For all the aforementioned analyses, we calculated social connectedness between a subject and each of their potential partners by combining pairwise grooming duration and spatial proximity into a dyadic composite sociality index (DSI) (35). DSI represents the relative rate at which a pair of individuals interact relative to the mean rate of interaction for all pairs of subjects in that given group and year. DSI can range from zero to infinity, with zero representing dyads that never interact and higher values representing dyads that spend more time interacting.

#### Prediction 4a

We found that female macaques showed a substantial within-individual increase in their proportion of kin partners with age. Females more than doubled the proportion of kin they interacted with over an eight-year period (120% increase), even when accounting for the increasing availability of kin that occurs as females reproduce and contribute offspring to the group (26)(within-individual age: *β* = 0.12; 95% CI = 0.03, 0.19; Figure 5a,b; Table S8).

**Fig. 5.**
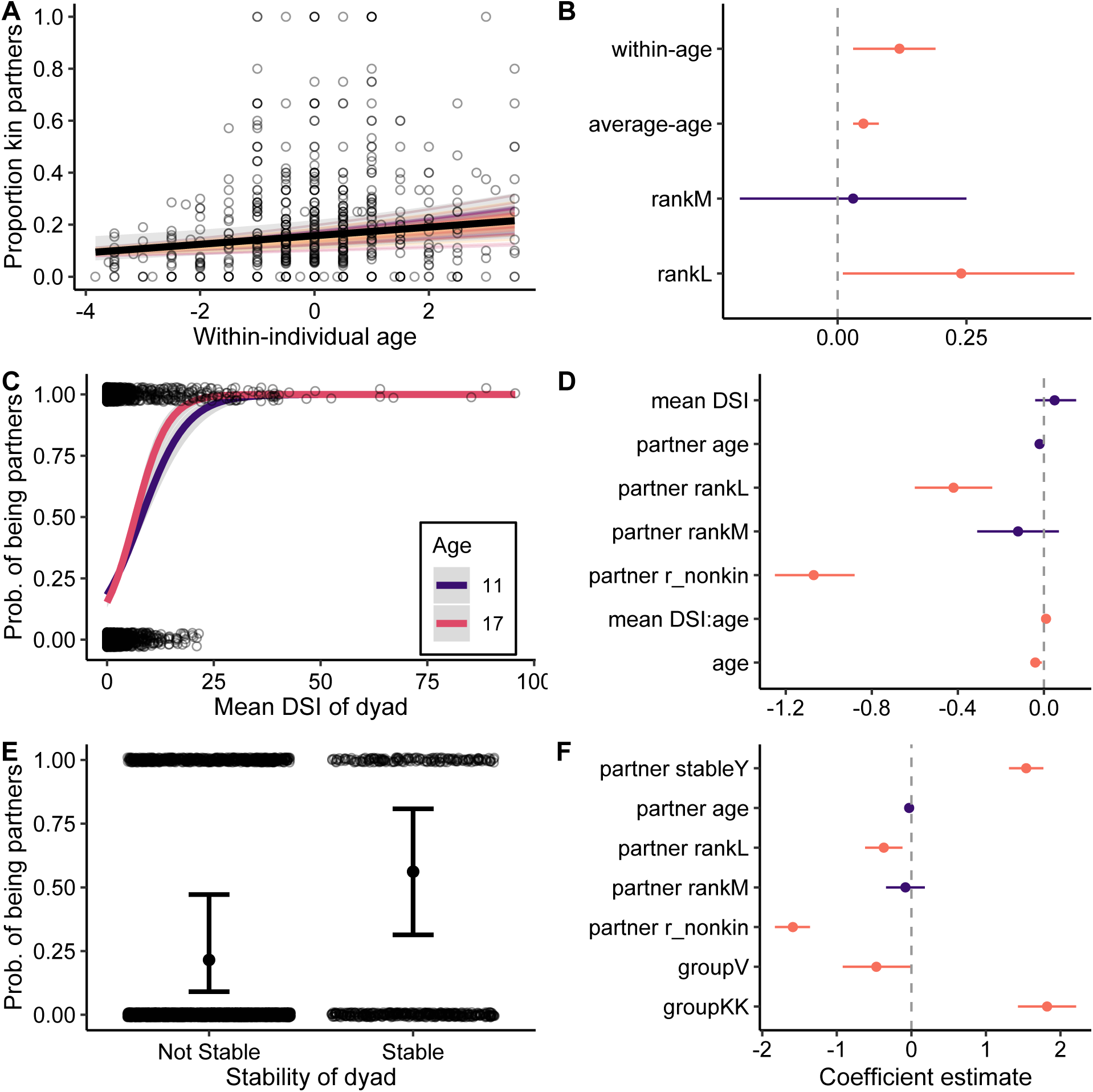
Individuals focus on important partners later in life. (**A**) Females show a within-individual increase in the proportion of kin partners with age. (**C** and **E**) Average dyadic composite sociality index (DSI) of a dyad and the stability of dyad positively predict the probability of being chosen as a partner by a female in later life (i.e. at the last point the female was observed in the dataset). Points represent raw data. Random slopes are shown using the thin coloured lines to illustrate the amount of inter-individual variation in a given social-aging pattern. Shaded gray ribbons and error bars indicate 95% confidence intervals around the predicted values. (**B, D** and **F**) Effect sizes and 95% credible intervals for all fixed effects for models that test the effect of (**B**) age on proportion of kin partners, (**D**) mean DSI of dyad on the probability of being partners with an older female and (**F**) stability of dyad on the probability of being partners with an older female. Instances where the 95% CI overlaps zero are colored in purple.

#### Prediction 4b & 4c

At a dyadic level, female macaques were also twice as likely to be partners with individuals to whom they had been strongly connected earlier in life (i.e. partners who had a high mean DSI value) and this effect was strongest in older females (mean DSI*age: *β* = 0.01; 95% CI = 0.01, 0.02; Figure 5c,d; Table S9). In older females (i.e. 17-years old) the most strongly connected individuals (those whose mean DSI values were greater than the mean value of 0.87) had on average a 39% chance of being a partner, while more poorly connected individuals (those with a mean DSI value < 0.87) only had on average a 17% chance of being a partner. Similarly, females were almost four times as likely to be partners with individuals with whom they had a stable social relationship earlier in life (i.e. partners with whom they had interacted for at least two consecutive years) (*β* = 1.54; 95% CI = 1.31, 1.77; Figure 5e,f, Table S10). Individuals who were not previously a stable partner had only a 21% chance of being a partner with females compared to individuals who were previously a stable partner and had a 56% chance of being a partner. This relationship was not age-dependent, in other words, we found no interaction between partner stability and female age. This result indicates that females are consistently pairing with stable partners regardless of their age and so this emphasis on stable partners may not be a behavior that is unique to older individuals. Regardless, both of these findings suggest that old females continue to emphasize important relationships despite narrowing their networks in later life.

### Little evidence of among-individual variation in the rate of aging

We found little evidence of among-individual variation in how individuals were changing their social behavior with age. In other words, there was little variation between individuals in the slope of the within-individual age term. In all of the aforementioned models the random slope term for individual ID explained between 0.16% and 0.64% of the variance in the model (see Tables S1-S8 Model A), while the random intercept term explained between 1.7% and 8.4% of the variance in the model. Thus there was much more variation in an individual’s average social behavior than how they changed their social behavior with age. The lack of variation among individuals in their slopes indicates that individuals change their behavior similarly throughout adulthood. In other words, an individual who ages from 12 to 18 is expected to show the same rate of change in social behavior as an individual who ages from 20 to 28.

### Age related changes in social behavior are not driven by selective disappearance

To test if selective disappearance was driving the observed age-dependent patterns of social behavior, we tested whether the within-individual age and average-age terms differed significantly from one another. We did this for all analyses by fitting a new model (Model B) where we replaced within-individual age in the original model (Model A; Equation 2) with age (Equation 3; see Methods for a more in-depth explanation). If the coefficient of average-age in Model B was “significantly” positive (or negative) it would indicate that individuals with low (or high) levels of the social behavior of interest selectively disappear from the population because the slopes of within-versus between-individual age differ significantly (15, 36). The 95% credible intervals for the average-age term in Model B always overlapped zero (see Tables S1-S8 Model B), meaning there was no evidence of selective disappearance driving age-related patterns in any of the models.

### Narrowing of social networks with increasing age is not influenced by deaths of social partners

We conducted a post-hoc analysis to account for the possibility that reductions in the number of partners with age might be driven by demographic changes, that is, by partners dying and not being replaced. Specifically, we asked whether the number of grooming partners and proximity partners in the current year (year t) was predicted by the number of social partners that died in the previous year (year t-1). The number of grooming partners in year t was not predicted by the number of partner deaths in year t-1 (*β* = 0.06; 95% CI = -0.04, 0.15; Table S11). The number of partner deaths in year t-1 did, however, have a significant positive effect on the number of proximity partners in year t (*β* = 0.12; 95% CI = 0.05, 0.18, Table S12). Therefore deaths of partners did not appear to be responsible for the narrowing of networks with age.

## Discussion

Increasing selectivity in social relationships is a commonly observed phenomenon in aging humans. Whether this pattern is unique to humans or characterizes other taxa must be addressed to fully understand the evolution of social aging. Here we show for the first time that within-individual changes in behavior across the lifespan can lead to increasing social selectivity in a non-human primate. We tested four key predictions of the social selectivity hypothesis and found that our results supported all of these predictions. We found that as females aged they reduced the size of their social networks and focused on kin, strongly-connected, and stable partners, all of which have been shown to provide important fitness benefits (21, 26). Females reduced the number of partners they approached as they aged, suggesting this narrowing was an active decision by the aging individual. Meanwhile, older females remained appealing social partners as they continued to be approached by similar numbers of partners and received similar amounts of grooming. Despite the reduction in partners with age, females remained actively engaged in the social sphere as they aged and continued to spend similar amounts of time giving grooming and in proximity to their social partners. Finally, the reduction in the number of partners with age was not driven by the deaths of social partners. In contrast, we observed that the more partners that died, the more females increased the number of partners they spent time in proximity to in the following year, perhaps as a means to compensate for the loss of those social relationships.

Our results provide the most comprehensive evidence to date for social selectivity in a non-human animal and ours is the first study to demonstrate that these patterns can result from within-individual changes in behavior. The distinction between within-individual and cohort-based change is critical because apparent declines in an individual’s network size or in other patterns of sociality can occur in the absence of behavioral changes across an individual’s lifetime. Because social selectivity is fundamentally a within-individual process, demonstrating behavioral changes across the lifespan is necessary evidence for this phenomenon. Here we demonstrate those within-individual changes and show that declines in sociality are not the result of selective disappearance or the deaths of social partners. While the use of cross-sectional data is an important means to approximate age-related changes in behavior when longitudinal data are not available, such results must be interpreted with caution. Studies that only use cross-sectional data or that cannot distinguish between-individual differences from within-individual changes may conflate social selectivity (i.e. behavioral plasticity) with population-level processes like selective disappearance. Alternatively, such studies might conclude there is no relationship between the social behavior of interest and age when there are actually two underlying associations of interest that counteract each other (15). Fundamentally, differentiating between within-individual changes in sociality with age and selective disappearance is important because it allows us to demonstrate that social behavior, just like other morphological, physiological or genomic traits, is a feature that can change across an individual’s lifetime. This perspective places sociality squarely within the larger aging phenotype and opens up the possibility of asking how and why these patterns of social aging have evolved and what their consequences are for other aspects of senescence.

Despite methodological differences, our results are consistent with many studies on human and other animals that find older individuals tend to have fewer social partners and to prioritize important relationships (3, 12, 19, 20, 32, 37-39). Similar to these studies, our findings indicate that female rhesus macaques demonstrate patterns of social aging that resemble the human social aging phenotype, suggesting that social selectivity may not solely arise from an awareness of limited time but may also be underpinned by other biological pathways. Many mammals face increased constraints and limitations as they age, including physiological changes as well as physical, energetic, and cognitive declines that might limit the capacity for, or alter the costs and benefits of, social interaction (16). Being more selective in partner choice and focusing on important or preferred partners with age might therefore reflect an adaptive response to these constraints. For example, rhesus macaques show declines in their immune function with age (25), which is a common phenomenon in mammals (40). This might select for withdrawal from social interactions to avoid competition and minimize the chance of negative encounters, or a reduction in network size to avoid contracting disease or illness. Similarly, declines in physical mobility or energetic capacity might select for individuals to be more discerning in which partners they spend their reduced effort and energy on.

Our findings that some age-related changes in sociality in female rhesus macaques were rank-dependent supports the idea that age-based changes in sociality may be an adaptive response to senescence. Specifically, lower ranking females decreased the number of partners they sat near and approached more strongly with age. In this population, older low-ranking females are more likely to be injured, which is strongly associated with increased mortality risk (41). Furthermore, low-ranking individuals exhibit greater increases in inflammatory cells with age (42). Thus, it is possible that older low-ranking females reduce their social integration more strongly with age to mitigate injury risk and the associated costs given their immune-compromised state. Old, low-ranking individuals have also been shown to avoid unpredictable social partners (43). This behavior might result from declines in information processing abilities, potentially as a result of elevated levels of adversity, which renders individuals unable to respond to social cues and adjust their behavior appropriately (43). Stronger declines in sociality with age among lower ranking individuals (32, 44) might therefore be an adaptive response to relatively greater cognitive and physiological constraints with age. It is also possible that lower-ranking individuals may experience more rapid senescence due to greater adversity (6, 45), which might accelerate declines in sociality with age.

Conclusively demonstrating that declines in sociality result from active selectivity with age remains challenging, and not only due to a lack of longitudinal data. Similar declines in sociality with age might occur as physical or mental deterioration inhibits an individual’s ability to interact with others or leads to reduced desirability of older individuals as social partners (16). Previous research has worked to disentangle these alternative hypotheses by showing, for instance, that Barbary macaques maintain an interest in vocal and visual social stimuli in later life (11, 46). Similar to our results, some studies have found that, despite interacting with fewer partners in later life, older individuals continue to receive the same or more affiliation from conspecifics, indicating that old individuals remain valuable social partners (11, 12, 26). In other cases, older individuals have been shown to engage in fewer energy-demanding activities (4, 46) or exhibit changes in space use with age (47) which invites the possibility that decreases in affiliation are not adaptive, but instead a direct consequence of deteriorating physical condition. Alternatively, social connectedness might be less strongly selected for at older ages, or declines in sociality could occur because enhanced social experience and skill among aged individuals free them from reliance on social capital to successfully navigate their environment (16). Previous research in this system has shown that the fitness benefits of social affiliation are stronger in prime-age females than in old-age females (26), in support of these possibilities.

Ultimately, distinguishing between alternative explanations for social aging remains an important avenue for future research. In this study, we have derived a clear set of predictions to test the social selectivity hypothesis. This approach increases confidence in the conclusion that social selectivity underlies the observed age-related declines in sociality in this macaque population and mitigates other potential explanations including energetic deficiencies, loss of social interest, reduced social desirability, loss of social partners, or selective disappearance. Nevertheless, future work on social aging will benefit from studies in which longitudinal changes in sociality with age can be measured alongside physical, energetic and cognitive changes to enable a fuller understanding of whether senescence precedes or follows changes in sociality across the lifespan. Perhaps even more critically, the fitness consequences of social aging will need to be explored to understand the adaptive nature of these changes. Future work should seek to assess age-specific selection on, and genetic architecture of, social traits to provide deeper insights into the evolution of age-dependency in sociality.

Given the well-established role that social integration plays in health and survival (6), understanding how social behavior changes with age and its associated fitness consequences will facilitate a deeper understanding of the mechanisms driving demographic aging under natural conditions. While social senescence is a topic that has been most extensively studied in primates (cf. 47, 48), there are many other group-living animals for whom social relationships are also critical for securing access to resources. As a result, age-related changes in sociality might play a pivotal role in life history tradeoffs between reproductive investment and somatic maintenance, thereby shaping senescence. Thus, there is an increasing need to incorporate social behavior into our broader understanding of the aging process across species if we are to better appreciate the forces shaping intra- and inter-individual variation in patterns of senescence.

## Methods

### Study Site and Population

We studied a population of free-ranging rhesus macaques on the island of Cayo Santiago off the southeastern coast of Puerto Rico. The animals are descendants of 409 Indian-origin rhesus macaques that were introduced to the island in the late 1930’s. The current population is maintained by the Caribbean Primate Research Centre (CPRC). All animals are food supplemented and provided with ad-libitum access to water. There are no predators on the island and there is no regular medical intervention for sick or wounded individuals, thus the major causes of death are illness and injury (49). Demographic data are collected up to five days per week by the CPRC staff and there is minimal dispersal from the island allowing for dates of birth and death for all individuals to be known to within a few days.

We began collecting behavioral data on individuals when they were considered to be “adults” (i.e. ≥ age 6), but here we focused on 204 females aged 10 years and older because we were specifically interesting in behavioral changes from prime adulthood into old age. Previous research on the Cayo Santiago population has shown that the median lifespan for females that survive to reproductive age is 15 years with a maximum lifespan of about 30 years (50). We focused on females that were alive between 2010-2017 and for whom we had detailed behavioral data, resulting in 563 macaque years of data, with an average of 2.8 years of data per individual (range: 1-8 years; Fig. 1). During this time period, we collected behavioral data from different study groups in different years (group F 2010-2017; group HH 2014; group KK 2015; group R 2015-2016; group S 2011; group V 2015-2016). We collected behavioral data using 10-min focal animal samples and recorded all behaviors continuously (51). We recorded the duration of grooming behavior along with the identities of the interactants and the direction of grooming. To establish spatial proximity, we conducted three scans at evenly spaced intervals during each focal sample and recorded the identities of all individuals within two meters of (but not in physical contact with) the study subject. We collected behavioral data between 07:30 and 14:00 and data collection was stratified to ensure equal sampling of individuals throughout the day and over the course of the year. For the purposes of this study, we included only interactions between adult females. We did not include interactions with males or juveniles of either sex as we wanted to avoid capturing changes in socio-sexual behavior with age, and the behavior of juveniles’ is commonly influenced by their lack of independence from their mother. We established dominance ranks for all females in a given year by using the direction and outcome of agonistic and submissive interactions (as per 21, 28). Rank was assigned as “high” (≥ 80% of other females dominated), “medium” (50%-79% of other females dominated) or “low” (≤ 49% of other females dominated).

### Quantifying social metrics

#### Prediction 1: Social networks will narrow with age

To quantify changes in the size of an individual’s social network with age, we explored changes in grooming degree and proximity degree. For each female in a given year, we calculated grooming degree as the number of unique individuals that a female gave grooming to or received grooming from and we calculated proximity degree as the number of unique individuals that a female was observed sitting in proximity to. We predicted to see declines in both grooming and proximity degree with age.

#### Prediction 2: Narrowing of networks will be driven by the aging individual

To determine if changes in network size were driven by the focal individual or by changes in the behavior of the social partners, we counted the unique number of individuals that the focal *approached* each year (approach outdegree) and the unique number of individuals that the focal was *approached by* each year (approach indegree). We predicted to see a decline in approach outdegree with age but no change in approach indegree with age.

#### Prediction 3: Individuals will show continued engagement and interest in the social world as they age

We quantified changes with age in the amount of grooming given (grooming outstrength) and received (grooming instrength) and the amount of time spent in proximity to other individuals (proximity strength). We calculated dyadic grooming outstrength and instrength as the total duration of grooming given and received by a subject (in seconds) divided by the total amount of time that both the subject and their partner were observed (in seconds). These dyadic measures were then summed across all pairs to give a total measure of individual grooming outstrength and instrength for each subject in each year. We calculated proximity strength in the same fashion. On a dyadic basis we first calculated the number of scans that pairs of individuals were in proximity to each other relative to the total number of scans done on both individuals. We then summed those dyadic measures across all pairs to give a total measure of individual proximity strength for each subject in each year. We predicted to see no change in the amount of time spent giving or receiving grooming or the amount of time spent in proximity with age.

#### Prediction 4: Individuals will focus on important relationships later in life

To test whether females focused in on important relationships with age we looked at three metrics: **(a)** whether females increased their proportion of kin partners with age, **(b)** whether older individuals were more likely to choose partners to whom they had previously been strongly connected, and **(c)** whether older individuals were more likely to choose partners with whom they had previously had stable social connections. We calculated social connectedness between a subject and each of their potential partners by combining grooming duration and spatial proximity into a dyadic composite sociality index (DSI) (35). Grooming and spatial proximity are two positively correlated (Pearson’s r = 0.37 in this study) affiliative social interactions that have been widely used to quantify the strength of dyadic bonds in primates (21, 27-30). We calculated total grooming duration between pairs of individuals and divided this by the total amount of time that both individuals were observed. This dyadic grooming rate was then divided by the mean grooming rate in that group/year. For proximity we again calculated the total number of scans that both individuals were in proximity and divided this by the total number of scans of both individuals. This dyadic proximity rate was then divided by the mean proximity rate in that group year. For each dyadic pair these standardized grooming and proximity rates were then summed and divided by the total number of behaviors (two) as per (35) to give the dyadic composite sociality index (DSI).

##### Prediction 4a

To assess changes in the proportion of kin partners with age we assigned all partners as either kin or non-kin using a cut off of r ≥ 0.125, as this is the level at which kin discrimination occurs for affiliative interactions in rhesus macaques from this population (52). Relatedness coefficients were calculated in the kinship2 package in R (version 1.8.5; 53) using the long-term pedigree maintained by the CPRC. We predicted to see an increase in the proportion of kin partners with age.

##### Prediction 4b

To assess whether older females were more likely to choose partners to whom they were strongly connected or had a stable relationship, we subset our data to only include the last year of data that we had for all subjects. For all subjects and their partners present in this “last-year” dataset (N = 11,050 dyads) we calculated the mean DSI using all previous years of data for the subject and their partner (range = 0-7 years, mean = 2 years of previous data on each dyad). Note that only subjects with at least two years of behavioral data could be included in this analysis (185 individuals), but we kept all potential partners in the last year dataset, including those that were observed in the group for the first time and so had a mean DSI of zero with the subject. It was important to account for the possibility that individuals appearing in the dataset for the first time might be chosen as partners by the subject despite not having previously been strongly connected. For this reason, it is possible for dyads to have zero years of previous data. It should be noted, however, that it is possible that some of these mean DSI values of zero may be false zeros. That is, because we only collected behavioral data on adults (i.e. ≥ age 6), it is possible that the focal individual and partner had a previously established relationship when the partner was a juvenile that was not captured in our dataset. However, the inclusion of these zeros only makes our analysis more conservative, and our results remained the same even when these potential partners with zero years of previous data were removed from the analysis. We predicted that individuals who were strongly connected to the subject earlier in life would be more likely to be partners with the subject in later life than individuals who were previously weakly connected to the subject.

##### Prediction 4c

For all subjects present in the last-year dataset we also assessed whether their partners were “stable” social partners (recorded as a categorical variable no/yes). Dyads were considered to be stable partners if they had a DSI > 0 for at least two consecutive years. Note that this means that only subjects with at least three years of behavioral data could be included in this analysis (113 individuals). For example, if a subject had two years of data, one year would be used in the last-year dataset, leaving only one year of previous data from which to calculate stability of social relationships. This means that by default all relationships with that subject would have been unstable as there would have been insufficient sampling time for stable social relationships to be established. As above, we kept all potential partners in the last-year dataset (N = 6,893 dyads). It was important to account for the possibility that individuals appearing in the dataset for the first time might be chosen as partners by the focal individual despite their lack of a stable relationship (again false zeros are possible because behavioral data collection began when individuals were adults). On average each dyad was observed for 2.7 years prior to the final year of data (range 0-7 years). Our results remained the same even when potential partners with only one or two years of data were removed from the analysis (i.e. partners with whom the subject did not have time to establish a stable relationship). We predicted that individuals who were stable partners with the subject earlier in life would be more likely to be partners with the subject in later life than individuals who were not stable partners previously.

### Statistical analyses

We used a suite of generalized linear mixed effects models in a Bayesian framework with different error structures and random effects to quantify changes in social behavior and partner preference with age.

#### Predictions 1-4a

For **Predictions 1 – 4a** all analyses were conducted at the level of the individual and we fitted two models for each analysis. In these analyses we were specifically interested in how social behavior changes across an individual’s lifetime, that is, in the within-subject effect of age. To separate the within-from the between-subject age components we used a within-subjects centering approach (as per 15, 54; see SI for details). Model A included within-individual age (to capture within-individual changes in behavior) and average age (to capture between-individual differences in behavior) as continuous fixed effects (see Equation 2 in the SI) as well as rank as a categorical fixed effect (see Tables S1-S8). We tested for an interaction between rank and within-individual age in all models to see if how social behavior changed across the lifespan was influenced by an individual’s social status, and removed the interaction when not significant. We included a random effect of group and year to account for variation in social behavior that might be due to differences between groups or years, and also included a random intercept term for individual ID, to account for repeated observations of the same females. We included within-individual age as a random slope term over individual ID, which allowed us to assess if there was variation in how individuals changed their social behavior with age. Although we did not expect non-linearities in the relationship between age and the response variable given that we were looking at changes in behavior from prime adulthood to old age, we nevertheless fitted a model with smoothing terms for within-individual age and average age and compared that model fit to the model with only linear age terms using leave-one-out cross-validation in the brms package (version 2.15.0; 55). Including smoothing terms in the model never improved the model fit and so was not considered further. For **Predictions 1-2** (grooming degree, proximity degree, approach outdegree and indegree) we fitted all models with a Poisson error distribution (log link). For **Prediction 3** (grooming outstrength and instrength, proximity strength) models were fitted using a zero-inflated beta regression error distribution (logit link). For **Prediction 4a** (proportion of kin partners) we fitted the model with a binomial error distribution (logit link) and also included proportion of kin in the group as an additional offset term to account for the increasing availability of kin with age.

Finally, we tested for selective disappearance by fitting a second model (Model B) for all the aforementioned models (see Tables S1-S8), which included age and average age (see Equation 3 in the SI) as well as the same fixed and random terms as Model A, but we did not include within-individual age as a random slope term over individual ID. In Model B the effect of age will be equivalent to the effect of within-individual age in Model A and the average age term now represents the difference between the between and within-subject effects. In cases where average age term is significant in Model B the between and within-individual slopes significantly differ, providing evidence for selective disappearance.

#### Predictions 4b and 4c

For **Predictions 4b and 4c** all analyses were conducted at the level of the dyad. We fitted whether or not individuals were partners (coded as 0/1) as the response variable in both models and used a Bernoulli error distribution (logit link). For **Prediction 4b** the predictor variable of interest was the mean DSI for the focal individual and their partner – calculated based on all previous years of interaction. This was included in the model as a continuous fixed effect. For **Prediction 4c** the predictor of interest was whether or not the focal individual and partner were stable social partners in previous years (no/yes). We included this in the model as a categorical fixed effect. We also included the partner’s age (continuous), the partner’s rank (categorical) and whether or not the partner was kin or non-kin (categorical) as fixed effects in both models (see Tables S9-S10). In each model we checked for an interaction between the predictor of interest (mean DSI and partner stability) and focal individual age to assess whether the likelihood of choosing a strong or stable partner was dependent on a female’s age. We removed the interaction term from the model when it was not significant. As above, we included group and year as random effects to account for variation in partner choice that might be due to differences between groups or years. Individual ID and partner ID were included as random effects in a multi-membership grouping term (56). This multi-membership grouping term accounts for the inherent multilevel structure of the data and allows each sample (dyad) to belong to more than one individual in a random effect at the same time.

#### Post-hoc analysis: Is network narrowing driven by deaths of social partners?

Finally, we conducted a post-hoc analysis to assess whether reductions in the number of grooming and proximity partners with age might be driven in part by deaths of social partners. To do this we fitted two models with a Poisson error distribution (log-link) where the response variables were number of grooming partners and number of proximity partners, respectively (see Tables S11-S12). In both models we fitted the number of partners that died in year t-1 as a continuous fixed effect (where a partner was any individual with a DSI > 0). We also included group as a categorical fixed effect since there were only three groups with at least two years of continuous data (R, V, and F). In all models we checked for an interaction between number of dead partners and female age to assess whether the effect of partner deaths was dependent on the age of the female. The interaction was not significant in either model and so was not considered further. As with all of the previous models we included year and individual ID as random effects.

We conducted all analyses using R version 4.1.0 (57) and fitted all models in the Bayesian software STAN (58) using the brms package (version 2.15.0; 55). All fixed effects were given weakly informative priors (see Supplementary Information for more details). We ran all models for 10,000 iterations across two chains with a warm-up period of 1,000 iterations. We assessed model convergence by examining traceplots to assess sampling mixing and by ensuring Rhat = 1. We considered estimates of fixed effects to be significantly different from zero when the 95% credible intervals of the posterior distribution did not overlap zero.

## Acknowledgments

We sincerely thank the Caribbean Primate Research Center for maintaining the Cayo population and for access to the study site. We are especially grateful to all the field technicians who have contributed to the long-term behavioral database over the years. We thank members of CRAB at the University of Exeter and members of the CBRU for thoughtful discussion during the manuscript’s development. This work was supported by the following grants from the National Institute of Health (NIH): grant nos R01-AG060931, R00-AG051764, R01-MH096875, R37-MH109728, R01-MH108627, R01-MH118203, U01MH121260, R01-NS123054 and the Kaufman Foundation: grant no KA2019-105548. Cayo Santiago Field Station is supported by the Office of Research Infrastructure Programs of the NIH (2P40OD012217).

## Data availability

All data and code associated with the analyses will be made publicly available on the Figshare Repository following acceptance of the manuscript.

