## Supplementary Materials for "Within-individual changes reveal increasing social selectivity with age in rhesus macaques"

**This PDF file includes:**

Supplementary text
Tables S1 to S12
Figures S1 to S4
SI References

### Supplementary Information Text

#### Separating within- from between-subject effects

In these analyses we were specifically interested in how social behavior changes across an individual's lifetime, that is, in the within-subject effect of age. To separate the within- from the between-subject age components we used a within-subjects centering approach (as per 1, 2). In a standard random effects linear regression, the estimated effect of a fixed term in the model includes both the within-subject effect and the between-subject effect, and is given by the following regression equation:

$$y_{ij} = \beta_0 + \beta_1 x_{ij} + u_{0j} + e_{0ij} \qquad \text{Equation 1}$$

where $x_{ij}$ is the $x$ value of measurement $i$ from subject $j$. $\beta_0$ is the intercept of the regression equation and $\beta_1$ is the slope showing the effect of predictor variable, $x$, on the response variable, $y$, for the $i^{th}$ measurement of subject $j$. The terms $u_{0j}$ and $e_{0ij}$ denote the random intercept and residual variance, respectively.

To isolate a within-individual effect, we separated age ($x_{ij}$) into two terms. The between-subject effect was obtained by taking the average age $\bar{x}$ of subject $j$, and was calculated by taking the mean age over all years that an individual was observed, resulting in one value of average age per individual $\bar{x}_j$. The within-subject effect (here called within-individual age) was calculated by subtracting average age from each age at which the individual was observed ($x_{ij} - \bar{x}_j$) resulting in many values per individual (one for each observation, centered around 0). This within-subject centering model is given by the regression equation:

$$y_{ij} = \beta_0 + \beta_W(x_{ij} - \bar{x}_j) + \beta_B \bar{x}_j + u_{0j} + e_{0ij} \qquad \text{Equation 2}$$

Here $\beta_W$ is the slope for the within-individual effect of age $(x_{ij} - \bar{x}_j)$ and $\beta_B$ is the slope for the between-individual effect of average age $\bar{x}_j$. This $\beta_B$ term therefore describes how the response variable (here some measure of social connectedness) is related to differences in age between individuals while $\beta_W$ describes how the response variable changes with age across an individual's lifetime, relative to their mean age.

We modified the equation above to account for the possibility that age-related changes in social behavior could be driven by selective disappearance of individuals with high or low measures of social connectedness. For instance, less social individuals may be more likely to die young because of poorer access to resources (3-5). To test for the presence of selective disappearance we transformed the equation above to test whether the slopes $\beta_B$ and $\beta_W$ differed significantly from one another (1, 2). The new formulation of the equation is as follows:

$$y_{ij} = \beta_0 + \beta_W x_{ij} + (\beta_B - \beta_W)\,\bar{x}_j + u_{0j} + e_{0ij} \qquad \text{Equation 3}$$

Here $\beta_W$ still represents the within-individual effect of age $x_{ij}$ and is equivalent to $\beta_W$ in Model 2 while the coefficient $(\beta_B - \beta_W)$ of average age $\bar{x}_j$ represents the difference between the between-individual and within-individual age terms. If this coefficient of average age is significantly positive it would indicate that individuals with low social connectedness (or with low values of the given response variable) disappear selectively from the population because the slopes of the within- versus between-subject age differ significantly (1, 2).

**Table S1.** Fixed and random effects from models looking at the effects of age on number of grooming partners (grooming degree). Model A is the within-individual centering model (based on Equation 2, see Methods), and Model B is the reformulation of this model (based on Equation 3, see Methods) to test for selective disappearance. Bolded terms indicate fixed effects where the 95% credible intervals did not overlap zero, providing evidence that those effects were significantly different from zero. Given that the 95% credible intervals for the average age term in Model B overlapped zero there was no evidence for selective disappearance.

We ran both models with the following weakly informative prior means and standard deviations (μ, σ): intercept (0, 2), within-age (0, 0.5), average-age (0, 0.5), rankL (0, 0.5), rankM (0, 0.5).

| Model | Effect | Group | Term | Estimate | Lower 95% CI | Upper 95% CI |
|---|---|---|---|---|---|---|
| **Model A** | Fixed Effects | | intercept | 2.01 | 1.52 | 2.52 |
| | | | **within-age** | **-0.06** | **-0.12** | **-0.01** |
| | | | **average-age** | **-0.04** | **-0.06** | **-0.02** |
| | | | **rankL** | **-0.42** | **-0.56** | **-0.28** |
| | | | **rankM** | **-0.27** | **-0.4** | **-0.12** |
| | Random Effects | Group | sd(intercept) | 0.38 | 0.15 | 0.91 |
| | | Year | sd(intercept) | 0.32 | 0.17 | 0.62 |
| | | Individual.ID | sd(intercept) | 0.22 | 0.14 | 0.30 |
| | | | sd(within-age) | 0.03 | 0.00 | 0.08 |
| | | | cor(intercept,within-age) | 0.10 | -0.92 | 0.95 |
| **Model B** | Fixed Effects | | intercept | 2.01 | 1.51 | 2.54 |
| | | | **age** | **-0.06** | **-0.11** | **-0.01** |
| | | | average-age | 0.02 | -0.04 | 0.08 |
| | | | **rankL** | **-0.42** | **-0.55** | **-0.29** |
| | | | **rankM** | **-0.27** | **-0.40** | **-0.13** |
| | Random Effects | Group | sd(intercept) | 0.39 | 0.15 | 0.92 |
| | | Year | sd(intercept) | 0.32 | 0.17 | 0.63 |
| | | Individual.ID | sd(intercept) | 0.22 | 0.14 | 0.29 |

**Table S2.** Fixed and random effects from models looking at the effects of age on number of proximity partners (proximity degree). Model A is the within-individual centering model (based on Equation 2, see Methods), and Model B is the reformulation of this model (based on Equation 3, see Methods) to test for selective disappearance. Bolded terms indicate fixed effects where the 95% credible intervals did not overlap zero, providing evidence that those effects were significantly different from zero. Given that the 95% credible intervals for the average age term in Model B overlapped zero there was no evidence for selective disappearance.

We ran both models with the following weakly informative prior means and standard deviations (μ, σ): intercept (0, 3), within-age (0, 0.5), average-age (0, 0.5), rankL (0, 0.5), rankM (0, 0.5), within-age:rankL (0, 0.5), within-age:rankM (0, 0.5).

| Model | Effect | Group | Term | Estimate | Lower 95% CI | Upper 95% CI |
|---|---|---|---|---|---|---|
| **Model A** | Fixed Effects | | intercept | 2.85 | 2.17 | 3.61 |
| | | | within-age | -0.01 | -0.08 | 0.05 |
| | | | **average-age** | **-0.03** | **-0.04** | **-0.01** |
| | | | rankM | -0.10 | -0.21 | 0.01 |
| | | | **rankL** | **-0.29** | **-0.4** | **-0.18** |
| | | | **within-age:rankL** | **-0.07** | **-0.13** | **-0.01** |
| | | | within-age:rankM | -0.03 | -0.09 | 0.03 |
| | Random Effects | Group | sd(intercept) | 0.61 | 0.29 | 1.35 |
| | | Year | sd(intercept) | 0.57 | 0.31 | 1.08 |
| | | Individual.ID | sd(intercept) | 0.26 | 0.22 | 0.31 |
| | | | sd(within-age) | 0.05 | 0.02 | 0.08 |
| | | | cor(intercept,within-age) | 0.67 | 0.17 | 0.98 |
| **Model B** | Fixed Effects | | intercept | 2.56 | 1.83 | 3.36 |
| | | | age | -0.03 | -0.08 | 0.01 |
| | | | average-age | 0.03 | -0.02 | 0.08 |
| | | | rankM | 0.24 | -0.1 | 0.57 |
| | | | rankL | 0.10 | -0.24 | 0.44 |
| | | | **age:ordinal.rankL** | **-0.03** | **-0.05** | **-0.01** |
| | | | age:ordinal.rankM | -0.03 | -0.05 | 0.00 |
| | Random Effects | Group | sd(intercept) | 0.64 | 0.30 | 1.39 |
| | | Year | sd(intercept) | 0.57 | 0.32 | 1.10 |
| | | Individual.ID | sd(intercept) | 0.26 | 0.22 | 0.31 |

**Figure S1.** Effects of age on within-individual changes in number of grooming partners and number of proximity partners (results shown on raw age scale based on Model B, Tables S1-2).

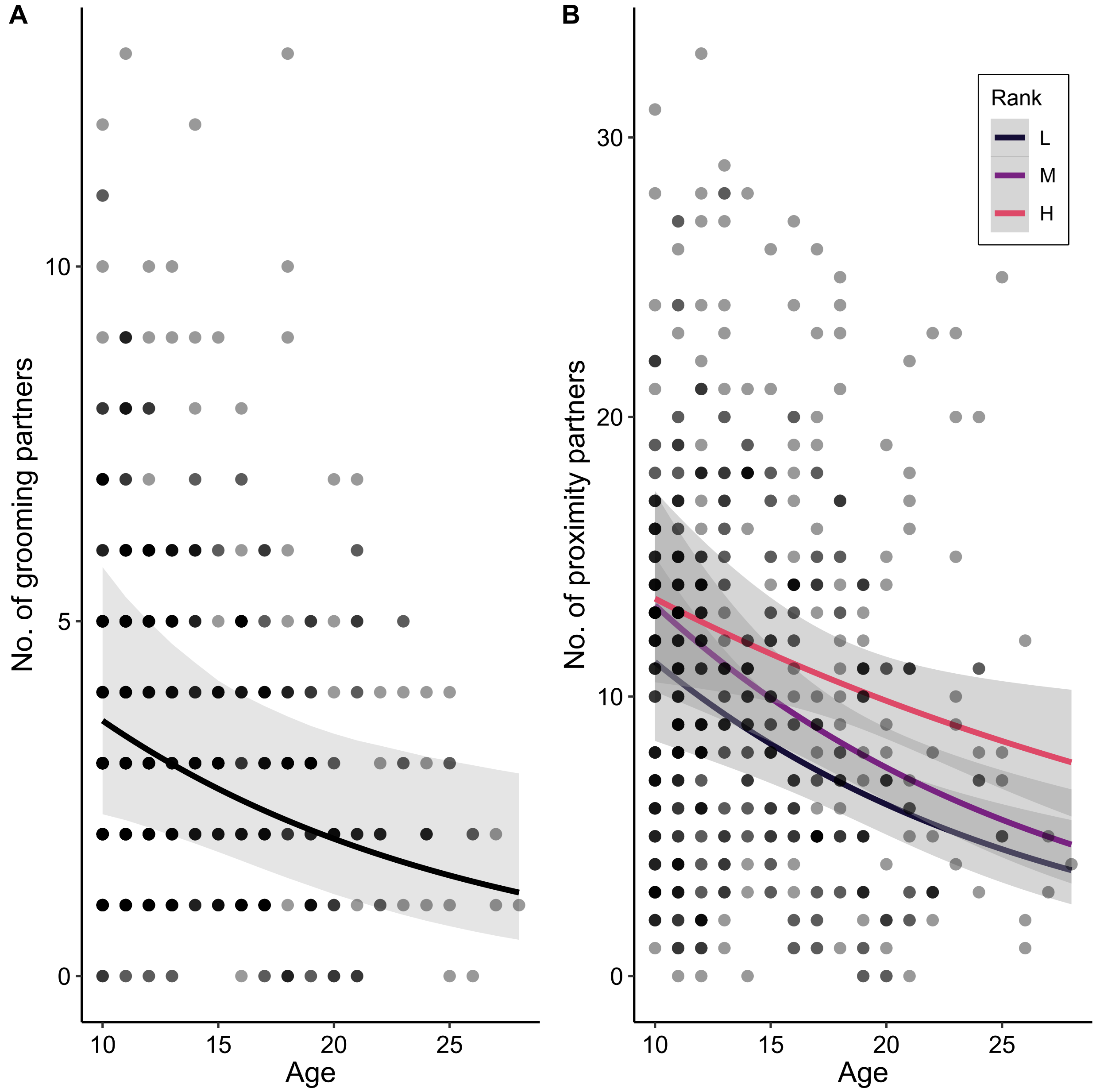

**Table S3.** Fixed and random effects from models looking at the effects of age on number of partners that the focal individual approached (approach outdegree). Model A is the within-individual centering model (based on Equation 2, see Methods), and Model B is the reformulation of this model (based on Equation 3, see Methods) to test for selective disappearance. Bolded terms indicate fixed effects where the 95% credible intervals did not overlap zero, providing evidence that those effects were significantly different from zero. Given that the 95% credible intervals for the average age term in Model B overlapped zero there was no evidence for selective disappearance.

We ran both models with the following weakly informative prior means and standard deviations (μ, σ): intercept (0, 3), within-age (0, 0.5), average-age (0, 0.5), rankL (0, 0.5), rankM (0, 0.5), within-age:rankL (0, 0.5), within-age:rankM (0, 0.5).

| Model | Effect | Group | Term | Estimate | Lower 95% CI | Upper 95% CI |
|---|---|---|---|---|---|---|
| **Model A** | Fixed Effects | | intercept | 2.73 | 2.08 | 3.37 |
| | | | within-age | -0.02 | -0.08 | 0.04 |
| | | | **average-age** | **-0.03** | **-0.04** | **-0.01** |
| | | | **rankL** | **-0.44** | **-0.55** | **-0.32** |
| | | | rankM | -0.09 | -0.2 | 0.02 |
| | | | **within-age:rankL** | **-0.08** | **-0.14** | **-0.01** |
| | | | within-age:rankM | -0.02 | -0.08 | 0.04 |
| | Random Effects | Group | sd(intercept) | 0.52 | 0.25 | 1.14 |
| | | Year | sd(intercept) | 0.56 | 0.31 | 1.05 |
| | | Individual.ID | sd(intercept) | 0.28 | 0.24 | 0.34 |
| | | | sd(within-age) | 0.06 | 0.03 | 0.08 |
| | | | cor(intercept,within-age) | 0.87 | 0.56 | 1.00 |
| **Model B** | Fixed Effects | | intercept | 2.54 | 1.87 | 3.21 |
| | | | age | -0.03 | -0.07 | 0.01 |
| | | | average-age | 0.02 | -0.03 | 0.07 |
| | | | rankL | -0.07 | -0.42 | 0.28 |
| | | | rankM | 0.11 | -0.21 | 0.44 |
| | | | **age:ordinal.rankL** | **-0.03** | **-0.05** | **-0.01** |
| | | | age:ordinal.rankM | -0.02 | -0.04 | 0.00 |
| | Random Effects | Group | sd(intercept) | 0.52 | 0.25 | 1.13 |
| | | Year | sd(intercept) | 0.57 | 0.31 | 1.07 |
| | | Individual.ID | sd(intercept) | 0.28 | 0.23 | 0.33 |

**Table S4.** Fixed and random effects from models looking at the effects of age on number of partners that the focal individual was approached by (approach indegree). Model A is the within-individual centering model (based on Equation 2, see Methods), and Model B is the reformulation of this model (based on Equation 3, see Methods) to test for selective disappearance. Bolded terms indicate fixed effects where the 95% credible intervals did not overlap zero, providing evidence that those effects were significantly different from zero. Given that the 95% credible intervals for the average age term in Model B overlapped zero there was no evidence for selective disappearance.

We ran both models with the following weakly informative prior means and standard deviations (μ, σ): intercept (0, 3), within-age (0, 0.5), average-age (0, 0.5), rankL (0, 0.5), rankM (0, 0.5).

| Model | Effect | Group | Term | Estimate | Lower 95% CI | Upper 95% CI |
|---|---|---|---|---|---|---|
| **Model A** | Fixed Effects | | intercept | 2.60 | 1.97 | 3.21 |
| | | | within-age | -0.02 | -0.07 | 0.02 |
| | | | **average-age** | **-0.03** | **-0.04** | **-0.01** |
| | | | rankL | -0.01 | -0.12 | 0.10 |
| | | | rankM | 0.05 | -0.05 | 0.15 |
| | Random Effects | Group | sd(intercept) | 0.48 | 0.22 | 1.07 |
| | | Year | sd(intercept) | 0.55 | 0.30 | 1.04 |
| | | Individual.ID | sd(intercept) | 0.27 | 0.22 | 0.31 |
| | | | sd(within-age) | 0.04 | 0.01 | 0.07 |
| | | | cor(intercept,within-age) | 0.70 | 0.16 | 0.98 |
| **Model B** | Fixed Effects | | intercept | 2.59 | 1.98 | 3.19 |
| | | | age | -0.02 | -0.06 | 0.02 |
| | | | average-age | -0.01 | -0.06 | 0.04 |
| | | | rankL | -0.04 | -0.15 | 0.06 |
| | | | rankM | 0.03 | -0.07 | 0.12 |
| | Random Effects | Group | sd(intercept) | 0.48 | 0.23 | 1.06 |
| | | Year | sd(intercept) | 0.54 | 0.3 | 1.03 |
| | | Individual.ID | sd(intercept) | 0.26 | 0.22 | 0.31 |

**Figure S2.** Effects of age on within-individual changes in number of partners approached and number of partners approached by with age (results shown on raw age scale based on Model B, Tables S3-4).

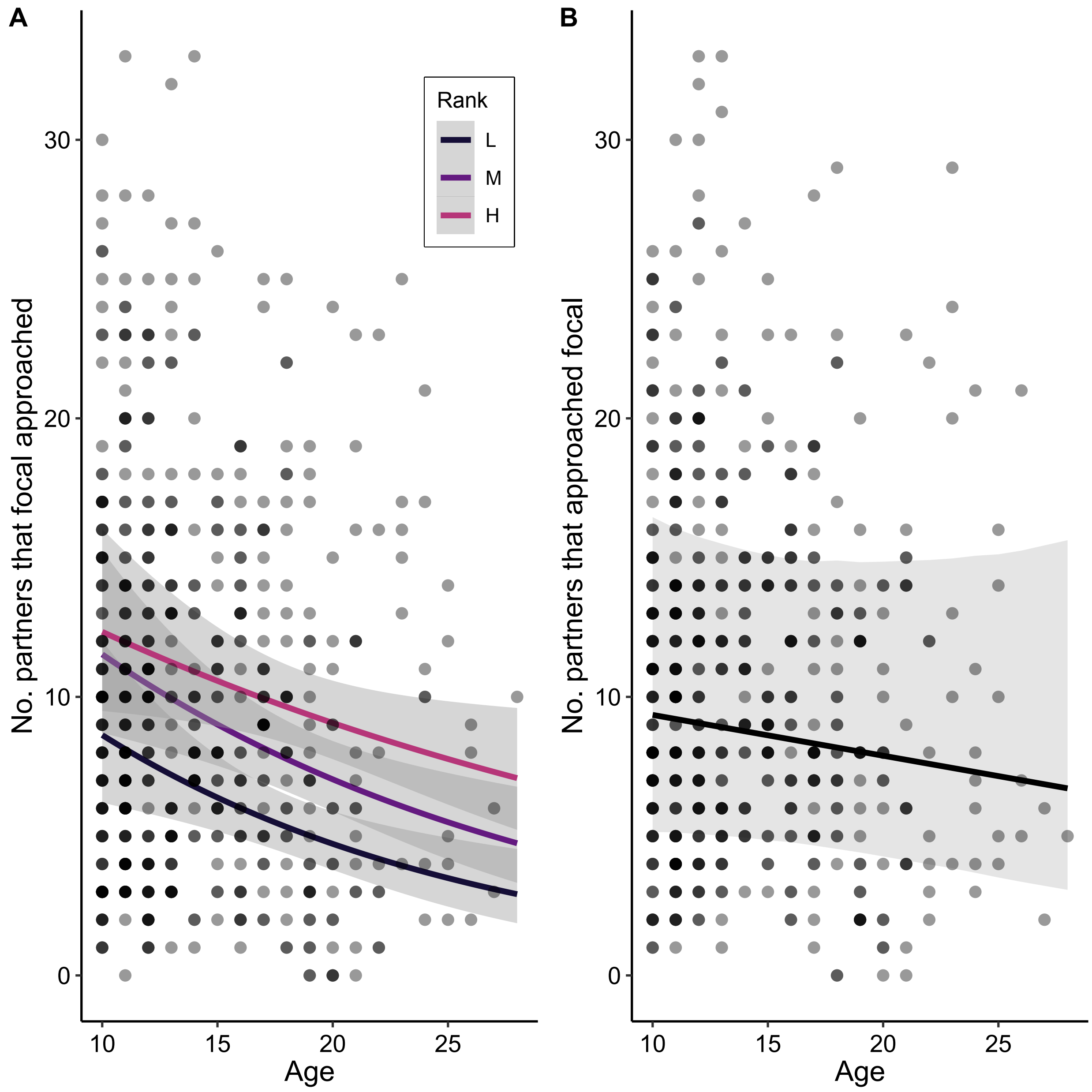

**Table S5.** Fixed and random effects from models looking at the effects of age on the amount of time spent giving grooming (grooming outstrength). Model A is the within-individual centering model (based on Equation 2, see Methods), and Model B is the reformulation of this model (based on Equation 3, see Methods) to test for selective disappearance. Bolded terms indicate fixed effects where the 95% credible intervals did not overlap zero, providing evidence that those effects were significantly different from zero. Given that the 95% credible intervals for the average age term in Model B overlapped zero there was no evidence for selective disappearance.

We ran both models with the following weakly informative prior means and standard deviations (μ, σ): intercept (0, 5), within-age (0, 0.5), average-age (0, 0.5), rankL (0, 0.5), rankM (0, 0.5).

| **Model** | **Effect** | **Group** | **Term** | **Estimate** | **Lower 95% CI** | **Upper 95% CI** |
|---|---|---|---|---|---|---|
| **Model A** | Fixed Effects | | intercept | -3.63 | -4.21 | -3.06 |
| | | | within-age | -0.05 | -0.12 | 0.03 |
| | | | average-age | -0.03 | -0.05 | 0.00 |
| | | | rankL | -0.11 | -0.30 | 0.08 |
| | | | rankM | -0.12 | -0.31 | 0.08 |
| | Random Effects | Group | sd(intercept) | 0.27 | 0.05 | 0.72 |
| | | Year | sd(intercept) | 0.39 | 0.18 | 0.80 |
| | | Individual.ID | sd(intercept) | 0.29 | 0.18 | 0.39 |
| | | | sd(within-age) | 0.07 | 0.01 | 0.15 |
| | | | cor(intercept,within-age) | -0.57 | -0.98 | 0.24 |
| **Model B** | Fixed Effects | | intercept | -3.62 | -4.16 | -3.06 |
| | | | age | -0.04 | -0.11 | 0.03 |
| | | | average-age | 0.02 | -0.06 | 0.10 |
| | | | rankL | -0.12 | -0.31 | 0.07 |
| | | | rankM | -0.12 | -0.32 | 0.07 |
| | Random Effects | Group | sd(intercept) | 0.25 | 0.05 | 0.65 |
| | | Year | sd(intercept) | 0.38 | 0.18 | 0.77 |
| | | Individual.ID | sd(intercept) | 0.28 | 0.17 | 0.39 |

**Table S6.** Fixed and random effects from models looking at the effects of age on the amount of time spent receiving grooming (grooming instrength). Model A is the within-individual centering model (based on Equation 2, see Methods), and Model B is the reformulation of this model (based on Equation 3, see Methods) to test for selective disappearance. Bolded terms indicate fixed effects where the 95% credible intervals did not overlap zero, providing evidence that those effects were significantly different from zero. Given that the 95% credible intervals for the average age term in Model B overlapped zero there was no evidence for selective disappearance.

We ran both models with the following weakly informative prior means and standard deviations ($\mu$, $\sigma$): intercept (0, 5), within-age (0, 0.5), average-age (0, 0.5), rankL (0, 0.5), rankM (0, 0.5).

| Model | Effect | Group | Term | Estimate | Lower 95% CI | Upper 95% CI |
|---|---|---|---|---|---|---|
| **Model A** | Fixed Effects | | intercept | -3.56 | -4.17 | -2.95 |
| | | | within-age | 0.02 | -0.05 | 0.09 |
| | | | average-age | -0.01 | -0.03 | 0.01 |
| | | | **rankL** | **-0.38** | **-0.54** | **-0.21** |
| | | | **rankM** | **-0.34** | **-0.52** | **-0.16** |
| | Random Effects | Group | sd(intercept) | 0.45 | 0.18 | 1.05 |
| | | Year | sd(intercept) | 0.37 | 0.17 | 0.75 |
| | | Individual.ID | sd(intercept) | 0.13 | 0.01 | 0.26 |
| | | | sd(within-age) | 0.05 | 0.00 | 0.12 |
| | | | cor(intercept,within-age) | -0.23 | -0.97 | 0.88 |
| **Model B** | Fixed Effects | | intercept | -3.55 | -4.16 | -2.93 |
| | | | age | 0.02 | -0.04 | 0.09 |
| | | | average-age | -0.03 | -0.11 | 0.04 |
| | | | **rankL** | **-0.38** | **-0.54** | **-0.21** |
| | | | **rankM** | **-0.35** | **-0.52** | **-0.17** |
| | Random Effects | Group | sd(intercept) | 0.45 | 0.18 | 1.07 |
| | | Year | sd(intercept) | 0.37 | 0.17 | 0.75 |
| | | Individual.ID | sd(intercept) | 0.13 | 0.01 | 0.26 |

**Table S7.** Fixed and random effects from models looking at the effects of age on the amount of time spent in proximity to other females (proximity strength). Model A is the within-individual centering model (based on Equation 2, see Methods), and Model B is the reformulation of this model (based on Equation 3, see Methods) to test for selective disappearance. Bolded terms indicate fixed effects where the 95% credible intervals did not overlap zero, providing evidence that those effects were significantly different from zero. Given that the 95% credible intervals for the average age term in Model B overlapped zero there was no evidence for selective disappearance.

We ran both models with the following weakly informative prior means and standard deviations ($\mu$, $\sigma$): intercept (0, 5), within-age (0, 0.5), average-age (0, 0.5), rankL (0, 0.5), rankM (0, 0.5).

| **Model** | **Effect** | **Group** | **Term** | **Estimate** | **Lower 95% CI** | **Upper 95% CI** |
|---|---|---|---|---|---|---|
| **Model A** | Fixed Effects | | intercept | -1.21 | -2.22 | -0.20 |
| | | | witihin-age | -0.02 | -0.08 | 0.05 |
| | | | average-age | -0.02 | -0.04 | 0.00 |
| | | | **rankL** | **-0.53** | **-0.68** | **-0.37** |
| | | | **rankM** | **-0.32** | **-0.47** | **-0.17** |
| | Random Effects | Group | sd(intercept) | 0.81 | 0.39 | 1.74 |
| | | Year | sd(intercept) | 0.86 | 0.48 | 1.61 |
| | | Individual.ID | sd(intercept) | 0.34 | 0.28 | 0.41 |
| | | | sd(witihin-age) | 0.08 | 0.01 | 0.20 |
| | | | cor(intercept,within-age) | -0.03 | -0.81 | 0.60 |
| **Model B** | Fixed Effects | | intercept | -1.21 | -2.23 | -0.18 |
| | | | age | -0.02 | -0.08 | 0.04 |
| | | | average-age | 0.00 | -0.07 | 0.08 |
| | | | **rankL** | **-0.51** | **-0.66** | **-0.37** |
| | | | **rankM** | **-0.35** | **-0.49** | **-0.21** |
| | Random Effects | Group | sd(intercept) | 0.81 | 0.40 | 1.76 |
| | | Year | sd(intercept) | 0.88 | 0.49 | 1.65 |
| | | Individual.ID | sd(intercept) | 0.34 | 0.27 | 0.40 |

**Figure S3.** Effects of age on within-individual changes in amount of grooming given and received and amount of time spent in proximity to other females (results shown on raw age scale based on Model B, Tables S5-7).

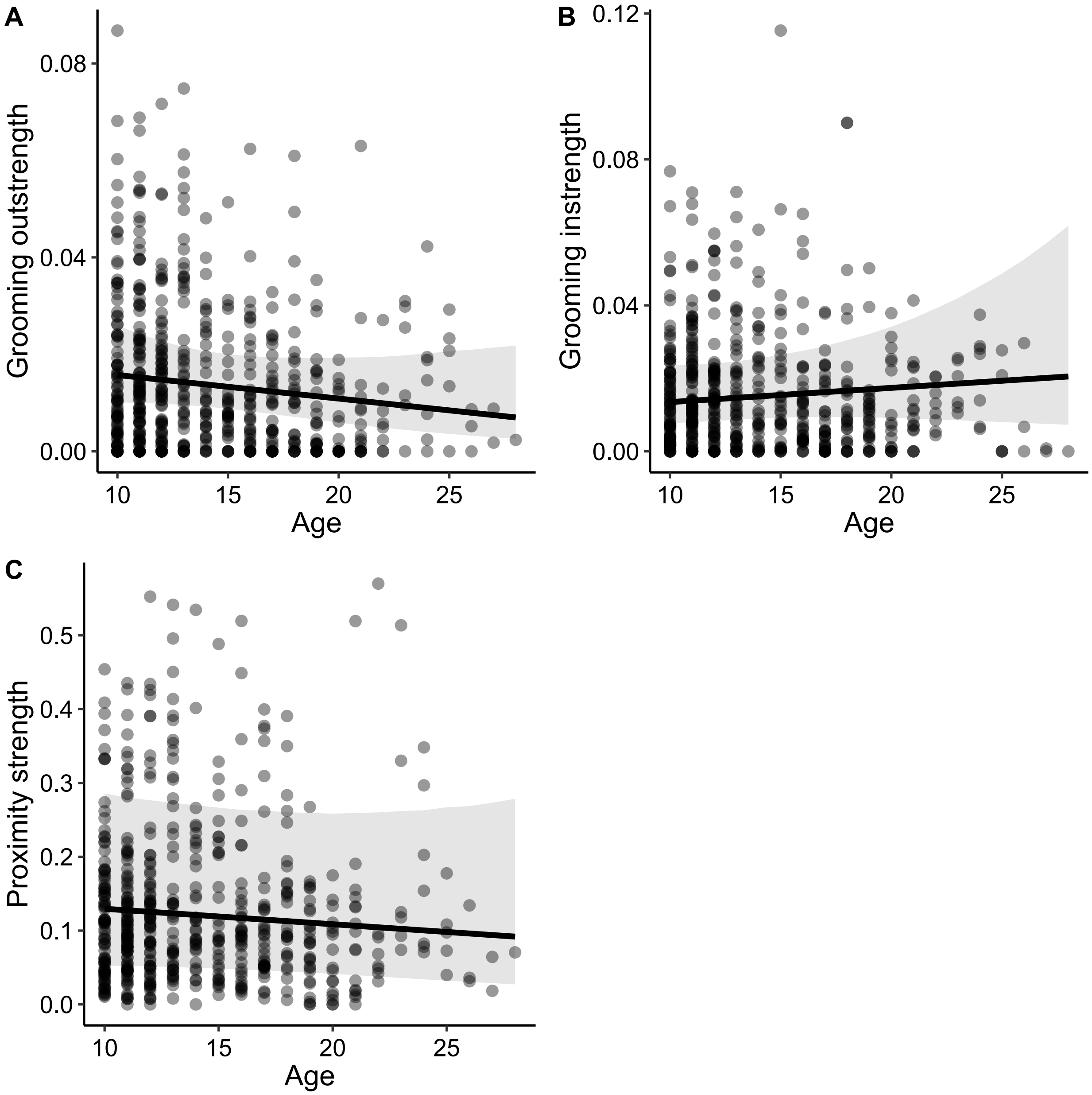

**Table S8.** Fixed and random effects from models looking at the effects of age on the proportion of kin partners. Model A is the within-individual centering model (based on Equation 2, see Methods), and Model B is the reformulation of this model (based on Equation 3, see Methods) to test for selective disappearance. Bolded terms indicate fixed effects where the 95% credible intervals did not overlap zero, providing evidence that those effects were significantly different from zero. Given that the 95% credible intervals for the average age term in Model B overlapped zero there was no evidence for selective disappearance.

We ran both models with the following weakly informative prior means and standard deviations (μ, σ): intercept (0, 5), within-age (0, 0.5), average-age (0, 0.5), rankL (0, 0.5), rankM (0, 0.5).

| Model | Effect | Group | Term | Estimate | Lower 95% CI | Upper 95% CI |
|---|---|---|---|---|---|---|
| **Model A** | Fixed Effects | | intercept | 0.06 | -0.72 | 0.80 |
| | | | **within-age** | **0.12** | **0.03** | **0.19** |
| | | | **average-age** | **0.05** | **0.03** | **0.08** |
| | | | **rankL** | **0.24** | **0.01** | **0.46** |
| | | | rankM | 0.03 | -0.19 | 0.25 |
| | Random Effects | Group | sd(intercept) | 0.65 | 0.28 | 1.44 |
| | | Year | sd(intercept) | 0.21 | 0.06 | 0.50 |
| | | Individual.ID | sd(intercept) | 0.28 | 0.14 | 0.41 |
| | | | sd(within-age) | 0.04 | 0.00 | 0.10 |
| | | | cor(intercept,within-age) | 0.32 | -0.85 | 0.98 |
| **Model B** | Fixed Effects | | intercept | 0.07 | -0.69 | 0.84 |
| | | | **age** | **0.12** | **0.03** | **0.19** |
| | | | average-age | -0.06 | -0.15 | 0.04 |
| | | | **rankL** | **0.24** | **0.02** | **0.46** |
| | | | rankM | 0.04 | -0.18 | 0.25 |
| | Random Effects | Group | sd(intercept) | 0.67 | 0.29 | 1.52 |
| | | Year | sd(intercept) | 0.21 | 0.06 | 0.49 |
| | | Individual.ID | sd(intercept) | 0.27 | 0.12 | 0.40 |

**Figure S4.** Effects of age on within-individual changes in proportion of kin partners (results shown on raw age scale based on Model B, Table S8).

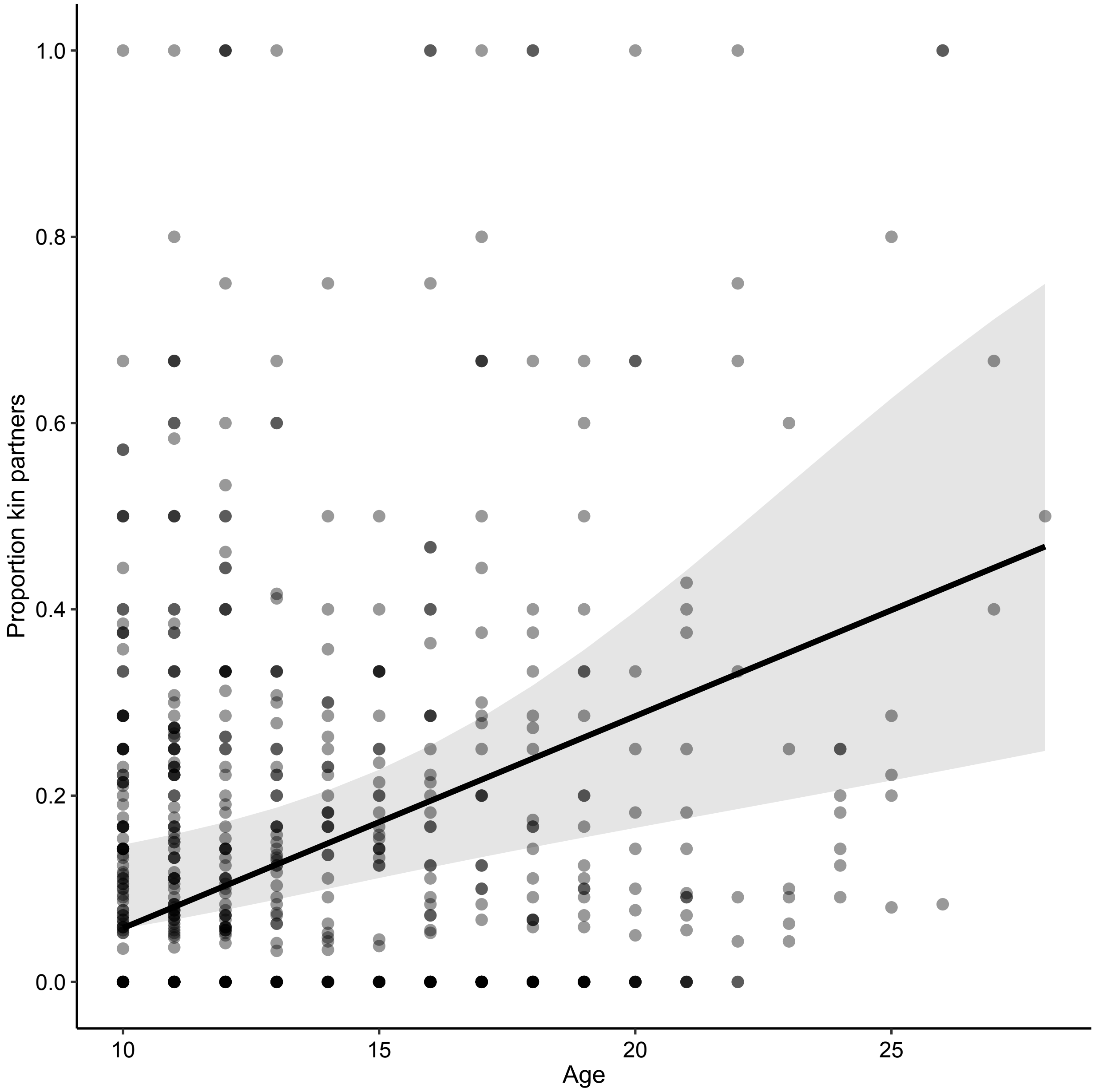

**Table S9.** Fixed and random effects from models looking at whether the mean dyadic sociality index (DSI) between a female and her partner predicts the probability of those individuals being partners later in the female's life. Bolded terms indicate fixed effects where the 95% credible intervals did not overlap zero, providing evidence that those effects were significantly different from zero. We ran the model with the following weakly informative prior means and standard deviations (μ, σ): intercept (0, 1), mean DSI (0, 1), partner age (0, 1), partner rankL (0, 1), partner rankM (0, 1), partner relatedness_nonkin (0, 1), mean DSI:age (0, 1).

| Effect | Group | Term | Estimate | Lower 95% CI | Upper 95% CI |
|---|---|---|---|---|---|
| **Fixed Effects** | | intercept | 0.67 | -0.51 | 2.05 |
| | | mean DSI | 0.05 | -0.04 | 0.15 |
| | | partner age | -0.02 | -0.04 | 0.00 |
| | | **partner rankL** | **-0.42** | **-0.60** | **-0.24** |
| | | partner rankM | -0.12 | -0.31 | 0.07 |
| | | **partner relatedness_nonkin** | **-1.07** | **-1.25** | **-0.88** |
| | | **age** | **-0.04** | **-0.06** | **-0.01** |
| | | **mean DSI:age** | **0.01** | **0.01** | **0.02** |
| **Random Effects** | Group | sd(intercept) | 1.31 | 0.58 | 2.95 |
| | Year | sd(intercept) | 0.91 | 0.44 | 1.84 |
| | Multi-membership | sd(intercept) | 0.87 | 0.74 | 1.01 |

**Table S10.** Fixed and random effects from models looking at whether the stability between a female and her partner (i.e. being partners for at least two consecutive years) predicts the probability of those individuals being partners later in the female's life. Bolded terms indicate fixed effects where the 95% credible intervals did not overlap zero, providing evidence that those effects were significantly different from zero. We ran the model with the following weakly informative prior means and standard deviations (μ, σ): intercept (0, 0.5), partner stableY (0, 1), partner age (0, 1), partner rankL (0, 1), partner rankM (0, 1), partner relatedness_nonkin (0, 1), groupV (0, 1), groupKK (0, 1).

| Effect | Group | Term | Estimate | Lower 95% CI | Upper 95% CI |
|---|---|---|---|---|---|
| Fixed Effects | | intercept | 0.70 | -0.41 | 1.93 |
| | | **partner stableY** | **1.54** | **1.31** | **1.77** |
| | | partner age | -0.03 | -0.05 | 0.00 |
| | | **partner rankL** | **-0.37** | **-0.62** | **-0.12** |
| | | partner rankM | -0.08 | -0.34 | 0.18 |
| | | **partner relatedness_nonkin** | **-1.59** | **-1.83** | **-1.36** |
| | | **groupV** | **-0.47** | **-0.92** | **-0.01** |
| | | **groupKK** | **1.82** | **1.43** | **2.21** |
| Random Effects | Year | sd(intercept) | 1.57 | 0.46 | 3.51 |
| | Multi-membership | sd(intercept) | 0.86 | 0.68 | 1.05 |

**Table S11.** Fixed and random effects from models looking at the effects of partner death in year t-1 on number of grooming partners in year t. Bolded terms indicate fixed effects where the 95% credible intervals did not overlap zero, providing evidence that those effects were significantly different from zero. We ran the model with the following weakly informative prior means and standard deviations (μ, σ): intercept (0, 1), no.partner.deaths (0, 0.5), groupR (0, 0.5), groupV (0, 0.5).

| Effect | Group | Term | Estimate | Lower 95% CI | Upper 95% CI |
|---|---|---|---|---|---|
| **Fixed Effects** | | intercept | 1.11 | 0.75 | 1.42 |
| | | no.partner.deaths | 0.06 | -0.04 | 0.15 |
| | | groupR | -0.15 | -0.39 | 0.08 |
| | | **groupV** | **-0.36** | **-0.63** | **-0.1** |
| **Random Effects** | Year | sd(intercept) | 0.4 | 0.2 | 0.83 |
| | Individual.ID | sd(intercept) | 0.3 | 0.22 | 0.4 |

**Table S12.** Fixed and random effects from models looking at the effects of partner death in year t-1 on number of proximity partners in year t. Bolded terms indicate fixed effects where the 95% credible intervals did not overlap zero, providing evidence that those effects were significantly different from zero. We ran the model with the following weakly informative prior means and standard deviations (μ, σ): intercept (0, 1), no.partner.deaths (0, 0.5), groupR (0, 0.5), groupV (0, 0.5).

| Effect | Group | Term | Estimate | Lower 95% CI | Upper 95% CI |
|---|---|---|---|---|---|
| **Fixed Effects** | | intercept | 1.92 | 1.25 | 2.41 |
| | | **no.partner.deaths** | **0.12** | **0.05** | **0.18** |
| | | groupR | 0.14 | -0.05 | 0.32 |
| | | **groupV** | **-0.68** | **-0.92** | **-0.45** |
| **Random Effects** | Year | sd(intercept) | 0.69 | 0.36 | 1.43 |
| | Individual.ID | sd(intercept) | 0.34 | 0.27 | 0.41 |
